# Comparative genomics of DH5α-inhibiting *Escherichia coli* isolates from human feces reveals common co-occurrence of bacteriocin genes with virulence factors and antibiotic resistance genes

**DOI:** 10.1101/2025.05.07.652706

**Authors:** Shuan Er, Yichen Ding, Linda Wei Lin Tan, Yik Ying Teo, Niranjan Nagarajan, Henning Seedorf

**Affiliations:** Temasek Life Sciences Laboratory, 1 Research Link, Singapore 117604, Singapore; Saw Swee Hock School of Public Health, National University of Singapore, 12 Science Drive 2, Singapore, 117549, Singapore; Department of Statistics and Data Science, National University of Singapore, Singapore, Singapore; Genome Institute of Singapore, A*STAR, Singapore, 138672, Singapore; Yong Loo Lin School of Medicine, National University of Singapore, Singapore, 117597, Singapore; Department of Biological Sciences, National University of Singapore, Singapore 117558, Singapore

## Abstract

The presence of multi-drug-resistant (MDR) bacteria in healthy individuals poses a significant public health concern, as these strains may contribute to or even facilitate the dissemination of antibiotic resistance genes (ARGs) and virulence factors. In this study, we investigated the genomic features of antimicrobial-producing *Escherichia coli* strains from the gut microbiota of healthy individuals in Singapore. Using a large-scale screening approach, we analyzed 3,107 *E. coli* isolates from 109 fecal samples for inhibitory activity against *E. coli* DH5α and performed whole-genome sequencing on 37 representative isolates.

Our findings reveal genetically diverse strains, with isolates belonging to five phylogroups (A, B1, B2, D, and F) and 23 unique sequence types (STs). Bacteriocin gene clusters were widespread, with colicins and microcins dominating the profiles. Notably, we identified a hcp-et3-4 gene cluster, encoding an effector linked to Type VI secretion system. Approximately 40% of the sequenced isolates were MDR, with resistance for up to eight antibiotic classes in one strain. Plasmids were the primary vehicles for ARG dissemination, but chromosomal resistance determinants were also detected. Additionally, over 55% of isolates were classified as potential extraintestinal pathogenic *E. coli* (ExPEC), raising concerns about their potential pathogenicity outside the intestinal tract.

Our study highlights the co-occurrence of bacteriocins, ARGs, and virulence genes in gut-residing *E. coli*, underscoring their potential role in shaping microbial dynamics and antibiotic resistance. While bacteriocin-producing strains show potential as probiotic alternatives, careful assessment of their safety and genetic stability is necessary for therapeutic applications.

**Importance:** This study provides genomic insights into the diverse repertoire of potential bacteriocin production in *E. coli* strains isolated from a cohort of healthy human subjects in Southeast Asia. Our findings suggest that the co-occurrence of bacteriocin genes with multiple antibiotic resistance and virulence genes is a common feature among non-clinical isolates. This genetic linkage could contribute to the increased abundance of multi-drug-resistant and virulent strains within individual hosts. Moreover, this co-occurrence may facilitate the host colonization with strains carrying antibiotic resistance and virulence determinants, potentially also contributing to the dissemination of resistant and pathogenic strains within the local population and beyond.

## Observation

The emergence of multi-drug resistant microorganisms is posing serious threats to global health care and calls for novel approaches to prevent the spread of infectious diseases. The rising prevalence of multi-drug resistant strains in healthy subjects is in this regard particularly worrisome as such individuals could contribute to dissemination of the resistance genes in populations and/or to pathogenic bacteria. In previous studies, we reported a high abundance of a multidrug-resistant *E. coli* strain, 94EC, in feces of a healthy subject that had not been treated with antibiotics prior to sampling (1, 2). This strain harbored resistance genes for seven different classes of antibiotics, including last-line antibiotics, such as tigecycline and colistin (3). It was noted that 94EC colonies displayed antimicrobial activity against other Enterobacteriaceae strains on agar plates (previously unpublished). The underlying causes for the antimicrobial activity and high abundance in the host remained unclear and whole-genome sequencing of the strain was therefore performed to analyze the genome for potential contributing features. Genome analysis revealed the presence of several bacteriocin gene clusters in the strain. Bacteriocins are known to have important roles in ecology and microbial population dynamics and have also been of interest for applications in biotechnological processes or as potential alternatives for antibiotic therapeutics. However, their co-occurrence with antibiotic resistance genes and their role for the spread of antibiotic resistance or in stabilizing the resistome are less well understood. Using samples from healthy subjects, we performed therefore a screen for *E. coli* strains that revealed antimicrobial activity against DH5α colonies (outlined in Figure 1). The aim was to determine how commonly antimicrobial activity co-occurs with antibiotic resistance and virulence genes in isolates from non-clinical subjects.

**Fig 1.**
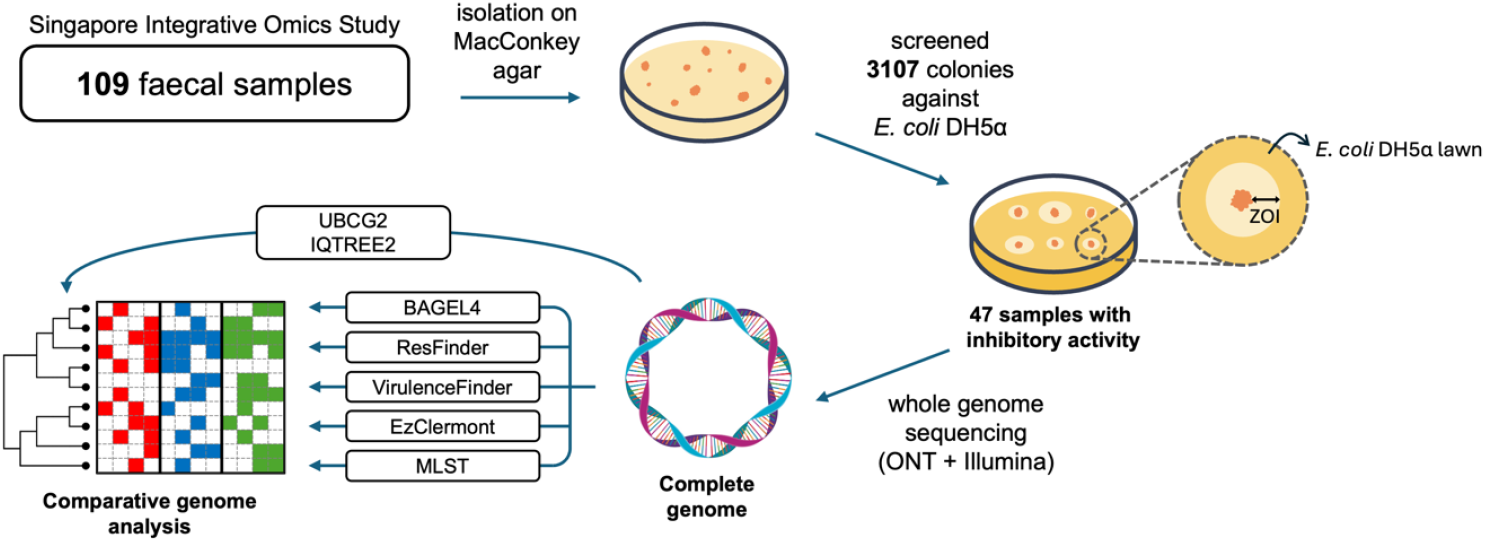
Schematic overview of workflow: fecal samples were screened on MacConkey agar, then against *E. coli* DH5*α* for inhibitory activity. Whole genome sequencing and comparative genomic analysis were carried out for positive isolates.

For the analysis, we performed a cultivation study on 109 fecal samples of healthy human individuals from the Singapore Integrative Omics Study cohort (see supporting information for details). Fecal samples were plated onto MacConkey agar to select for gram-negative bacteria. We then screened 3107 colonies against common laboratory *Escherichia coli* strain DH5*α* in a large-scale spot-on-lawn assay. One representative isolate was picked from each sample that was positive for inhibitory activity against *E. coli* DH5*α. E. coli* strain 94EC was included in this dataset due to its potential clinical significance. Comparative analysis was then performed for complete whole genomes derived from a total of 38 *E. coli* strains. Presence of bacteriocin, antibiotic resistance and virulence genes were predicted with BAGEL4, ResFinder v4.5.0 and VirulenceFinder v2.0.5 (4-8). Phylogroups and sequence types were determined using EzClermont and MLST v2.0.9 (9, 10). Core genes were extracted and aligned using UBCG2 and the maximum likelihood phylogenetic tree was inferred using IQ-TREE v2.3.6 (11, 12). The *E. coli* isolates (Figure 2) were predicted to belong to five distinct phylogroups. The most dominant phylogroups are A (15/38, 39.5%) and D (8/39, 21.1%), followed by B1 (6/38, 15.8%), B2 (5/38, 13.2%) then F (4/38, 10.5%). All 38 isolates were assigned to 23 unique sequence types (STs) and one unassigned (D96). Of note, ST10 was the most prevalent (8 isolates), followed by ST69 (4 isolates); among the remaining STs, five were doubletons and 17 were singletons (including one unassigned). The core genome phylogenetic tree illustrates the distribution of phylogroups and STs. Specifically, phylogroups B2 and F fall into a distinct group (Group I) comprising nine isolates belonging to seven unique STs while A, B1 and D make up another group (Group II) consisting of 29 isolates belonging to 17 unique STs.

**Fig 2.**
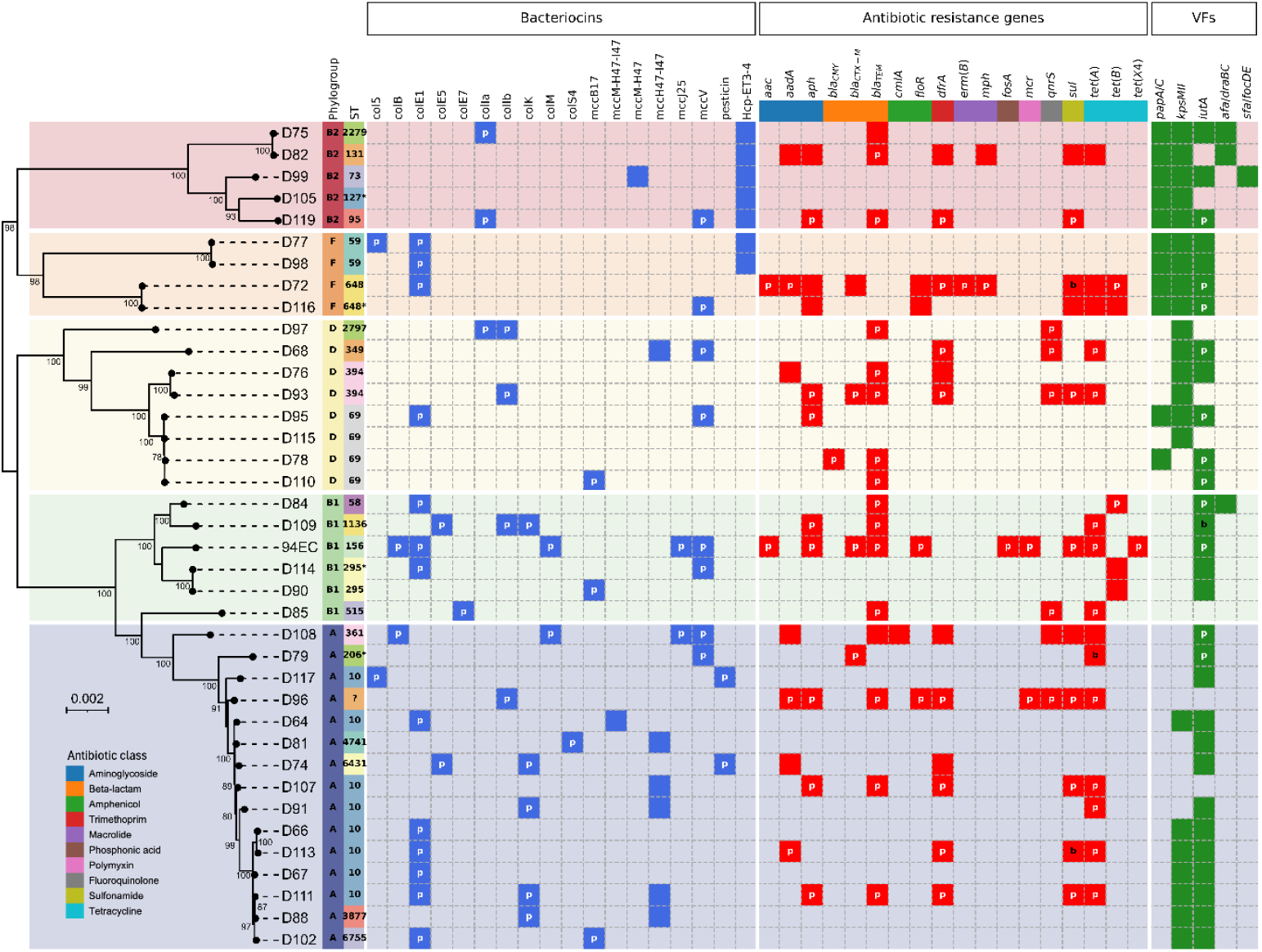
Heatmap illustrating the distribution of genomic factors among the *E. coli* isolates. Presence of bacteriocin, antibiotic resistance and virulence gene clusters are marked by colored tiles blue, red and green, respectively. Non-solid tiles indicate absence of genes. Genomic location of the genes is represented by letters: none – chromosome, p – plasmid, b – both chromosome and plasmid. Maximum likelihood phylogenetic tree was constructed using a concatenated alignment of 81 core genes supported by 1000 ultrafast bootstrap replicates. Only bootstrap values ≥ 70% are shown. Scale bar: 0.002 nucleotide substitutions per site.

Bacteriocin gene cluster prediction reveals a vast spread of bacteriocin types. A total of 17 bacteriocin variants were identified, of which, ten are colicins, six are microcins and one pesticin. Our findings are concordant with the fact that colicins are typically plasmid-encoded while microcins can be either plasmid- or chromosomally encoded (13). Of the 38 isolates, seven harbored 3-5 bacteriocin variants, twenty-eight harbored 1-2 variants while three (found exclusively in phylogroup D) did not possess any known bacteriocins despite producing inhibitory activity against *E. coli* DH5*α* growth. ColE1 was the most frequently occurring bacteriocin, identified in 13 out of 38 isolates (34.2%). Majority of ColE1 carriers belonged to phylogroups A, B1 and F, and were absent from phylogroup B2 and only one of nine phylogroup D isolates possessed this bacteriocin.

Interestingly, we identified a chromosomal *hcp-et3-4* gene cluster in seven isolates. Although the Hcp-ET3-4 gene product is an effector of the type VI secretion system (T6SS), it contains the Pyocin S (ET3) and Colicin-DNase (ET4) bacteriocin domains (14), hence, we classified this gene cluster as a bacteriocin variant for the scope of this study. We found that all isolates harboring *hcp-et3-4* were restricted to Group I – five phylogroup B2 isolates and two phylogroup F isolates. Among the other chromosomally-encoded gene clusters, we also identified siderophore-microcins MccM-H47-I47, MccM-H47 and MccM-H47-I47. Of note, the MccH47-I47 cluster was the most prevalent and exclusively carried by Group II isolates – six in phylogroup A isolates and one in phylogroup D. The other two clusters were each present in a single isolate. Our findings contrast with a previous study conducted on clinical samples which reported a higher prevalence in phylogroup B2 *E. coli* (15). A possible reason for this observation could be the difference in sample sources – isolates in this study were derived from fecal samples of healthy subjects whereas Š majs et al. analyzed isolates from urinary tract infections (UTIs). Phylogroup B2 *E. coli* are often associated with UTIs, and MccH47 has been suggested to be an important determinant for facilitating colonization and subsequent emergence of phylogroup B2 strains from the intestinal reservoir (16, 17). Hence, this may explain the higher incidence of MccH47 observed in clinical UTI samples compared to our dataset.

The ARG profiles of the isolates can provide valuable insights into the gut antibiotic resistance landscape of healthy individuals in Singapore. Prediction of ARG clusters identified resistance to 12 unique classes of antibiotics – most prevalent being tetracycline and beta-lactam (each 18/38, 47.4%), followed by aminoglycoside (15/38, 39.5%), trimethoprim (12/38, 31.6%), sulfonamide (11/38, 28.9%) and fluoroquinolone (6/38, 15.8%). About one-third of all isolates were predicted to not carry any known ARGs.

Strains are considered multidrug resistant (MDR) if they are resistant to at least three distinct classes of antibiotics (18). Our analysis revealed that nearly 40% of isolates (15/38) are MDR. Most prominently, D96 was predicted to be resistant to 8 classes; D72, D108, 94EC resistant to 7 classes; D82, D93 resistant to 6 classes. A majority of ARGs were located on plasmids (68.7%); 28.4% were located on chromosome only and 2.9% were found on both. Interestingly, we observed that eight isolates have most, if not all, ARGs located on the chromosome, out of which, half of them are MDR. This presents an opportunity for investigating the evolution of the intrinsic (chromosomal) resistome which is necessary for predicting the likelihood of emergence of antibiotic resistance in bacterial populations (19).

In addition to bacteriocin and ARG profiling, the diversity of virulence factors (VFs) among these isolates was also elucidated. On average, Group I isolates carried substantially more VFs than Group II isolates (Group I: 33.2 ± 6.9 VFs, Group II: 22.9 ± 5.7 VFs. Phylogroups B2 and F (Group I) isolates have frequently been implicated as extraintestinal pathogenic *E. coli* (ExPEC) in humans, capable of causing infections outside of the gastrointestinal tract. In our dataset, we found that 55.3% (21/38) of the isolates were considered ExPEC (≥2 ExPEC VF markers (20)). A further breakdown revealed that two isolates (D75 and D99) carried four ExPEC markers, seven isolates carried three ExPEC markers and 12 isolates harbored two ExPEC markers. Indeed, it was observed that all Group I isolates were ExPEC strains and possessed ≥3 ExPEC markers (except D105) whereas slightly less than half of Group II isolates were ExPEC and all of them carried ≤2 markers, except D95 (phylogroup D) which carried 3 markers. ExPEC strains are of clinical significance as they can cause a myriad of non-intestinal infections in the human body as well as have the ability to transmit resistance genes to other pathogenic bacteria. Our study showed a relatively high proportion of ExPEC strains present in the feces of healthy human subjects, furthermore, about one-third of these are MDR, thereby raising concerns of potential problematic infections by these opportunistic pathogens.

Here, we show evidence for a diverse profile of DH5*α*-inhibiting *E. coli* in a healthy Singaporean cohort. These strains were made up of at least 23 unique STs belonging to five phylogroups and most were bacteriocinogenic. Furthermore, a large proportion of them were MDR and ExPEC strains. Approximately 70% of detected ARGs were located on plasmids which may then act as molecular vehicles to spread resistance genes to highly virulent pathogens, thus, posing a threat to treatment by conventional antibiotics. Nonetheless, bacteriocinogenic *E. coli* from our dataset may also show potential as promising therapeutic alternatives (probiotics) to antibiotics. A key advantage lies in the specificity of the bacteriocins they produce which can selectively target pathogenic bacteria (typically close relatives of the bacteriocin producer), thus, the narrow antimicrobial spectrum minimizes disruption to the surrounding microbiota. For instance, Mortzfeld et al. showed that MccI47 exhibits specific inhibitory activity against *Enterobacteriaceae* strains and its potency is comparable to commonly prescribed antimicrobials (21). The targeted antimicrobial activity of bacteriocins, combined with their origin as commensal gut microbes, makes the isolates in our study suitable as probiotic candidates. However, safety and efficacy must be ensured by selecting isolates devoid of ARGs and can also be further optimized by genetic engineering to remove undesirable genomic elements.

## References

1. Ding Y, Er S, Tan A, Gounot J-S, Saw W-Y, Tan LWL, Teo YY, Nagarajan N, Seedorf H. 2024. Comparison of tet(X4)-containing contigs assembled from metagenomic sequencing data with plasmid sequences of isolates from a cohort of healthy subjects. Microbiology Spectrum 12:e03969–23.

2. Ding Y, Saw W-Y, Tan LWL, Moong DKN, Nagarajan N, Teo YY, Seedorf H. 2020. Emergence of tigecycline- and eravacycline-resistant Tet(X4)-producing Enterobacteriaceae in the gut microbiota of healthy Singaporeans. Journal of Antimicrobial Chemotherapy 75:3480–3484.

3. Ding Y, Saw W-Y, Tan LWL, Moong DKN, Nagarajan N, Teo YY, Seedorf H. 2021. Extended-Spectrum β-Lactamase-Producing and mcr-1-Positive Escherichia coli from the Gut Microbiota of Healthy Singaporeans. Applied and Environmental Microbiology 87:e00488–21.

4. van Heel AJ, de Jong A, Song C, Viel JH, Kok J, Kuipers OP. 2018. BAGEL4: a user-friendly web server to thoroughly mine RiPPs and bacteriocins. Nucleic Acids Research 46:W278–W281.

5. Bortolaia V, Kaas RS, Ruppe E, Roberts MC, Schwarz S, Cattoir V, Philippon A, Allesoe RL, Rebelo AR, Florensa AF, Fagelhauer L, Chakraborty T, Neumann B, Werner G, Bender JK, Stingl K, Nguyen M, Coppens J, Xavier BB, Malhotra-Kumar S, Westh H, Pinholt M, Anjum MF, Duggett NA, Kempf I, Nykäsenoja S, Olkkola S, Wieczorek K, Amaro A, Clemente L, Mossong J, Losch S, Ragimbeau C, Lund O, Aarestrup FM. 2020. ResFinder 4.0 for predictions of phenotypes from genotypes. Journal of Antimicrobial Chemotherapy 75:3491–3500.

6. Camacho C, Coulouris G, Avagyan V, Ma N, Papadopoulos J, Bealer K, Madden TL. 2009. BLAST+: architecture and applications. BMC Bioinformatics 10:421.

7. Joensen KG, Scheutz F, Lund O, Hasman H, Kaas RS, Nielsen EM, Aarestrup FM. 2014. Real-Time Whole-Genome Sequencing for Routine Typing, Surveillance, and Outbreak Detection of Verotoxigenic Escherichia coli. Journal of Clinical Microbiology 52:1501–1510.

8. Tetzschner AMM, Johnson JR, Johnston BD, Lund O, Scheutz F. 2020. In Silico Genotyping of Escherichia coli Isolates for Extraintestinal Virulence Genes by Use of Whole-Genome Sequencing Data. Journal of Clinical Microbiology 58:10.1128/jcm.01269-20.

9. Larsen MV, Cosentino S, Rasmussen S, Friis C, Hasman H, Marvig RL, Jelsbak L, Sicheritz-Pontén T, Ussery DW, Aarestrup FM, Lund O. 2012. Multilocus Sequence Typing of Total-Genome-Sequenced Bacteria. Journal of Clinical Microbiology 50:1355–1361.

10. Waters NR, Abram F, Brennan F, Holmes A, Pritchard L. 2020. Easy phylotyping of Escherichia coli via the EzClermont web app and command-line tool. Access Microbiology 2.

11. Kim J, Na S-I, Kim D, Chun J. 2021. UBCG2: Up-to-date bacterial core genes and pipeline for phylogenomic analysis. Journal of Microbiology 59:609–615.

12. Minh BQ, Schmidt HA, Chernomor O, Schrempf D, Woodhams MD, von Haeseler A, Lanfear R. 2020. IQ-TREE 2: New Models and Efficient Methods for Phylogenetic Inference in the Genomic Era. Molecular Biology and Evolution 37:1530–1534.

13. Arbulu S, Kjos M. 2024. Revisiting the Multifaceted Roles of Bacteriocins : The Multifaceted Roles of Bacteriocins. Microb Ecol 87:41.

14. Ma J, Pan Z, Huang J, Sun M, Lu C, Yao H. 2017. The Hcp proteins fused with diverse extended-toxin domains represent a novel pattern of antibacterial effectors in type VI secretion systems. Virulence 8:1189–1202.

15. Šmajs D, Micenková L, Šmarda J, Vrba M, Ševčíková A, Vališová Z, Woznicová V. 2010. Bacteriocin synthesis in uropathogenic and commensal Escherichia coli: colicin E1 is a potential virulence factor. BMC Microbiology 10:288.

16. Hogins J, Xuan Z, Zimmern PE, Reitzer L. 2023. The distinct transcriptome of virulence-associated phylogenetic group B2 Escherichia coli. Microbiology Spectrum 11:e02085–23.

17. Massip C, Oswald E. 2020. Siderophore-Microcins in Escherichia coli: Determinants of Digestive Colonization, the First Step Toward Virulence. Front Cell Infect Microbiol 10:381.

18. Magiorakos AP, Srinivasan A, Carey RB, Carmeli Y, Falagas ME, Giske CG, Harbarth S, Hindler JF, Kahlmeter G, Olsson-Liljequist B, Paterson DL, Rice LB, Stelling J, Struelens MJ, Vatopoulos A, Weber JT, Monnet DL. 2012. Multidrug-resistant, extensively drug-resistant and pandrug-resistant bacteria: an international expert proposal for interim standard definitions for acquired resistance. Clinical Microbiology and Infection 18:268–281.

19. Martínez JL, Baquero F, Andersson DI. 2007. Predicting antibiotic resistance. Nature Reviews Microbiology 5:958–965.

20. Johnson JR, Murray AC, Gajewski A, Sullivan M, Snippes P, Kuskowski MA, Smith KE. 2003. Isolation and molecular characterization of nalidixic acid-resistant extraintestinal pathogenic Escherichia coli from retail chicken products. Antimicrob Agents Chemother 47:2161–8.

21. Chakraborty A, Saralaya V, Adhikari P, Shenoy S, Baliga S, Hegde A. 2015. Characterization of Escherichia coli Phylogenetic Groups Associated with Extraintestinal Infections in South Indian Population. Ann Med Health Sci Res 5:241–6.

